## Supplementary Materials for "Flexing the principal gradient of the cerebral cortex to suit changing semantic task demands"

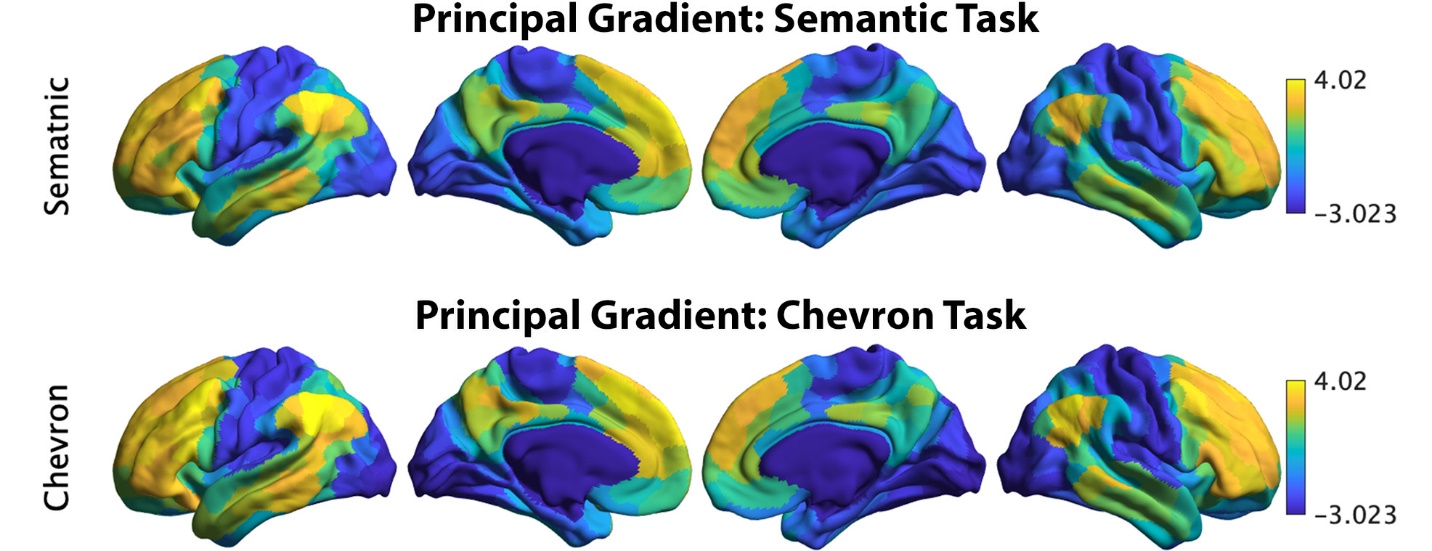

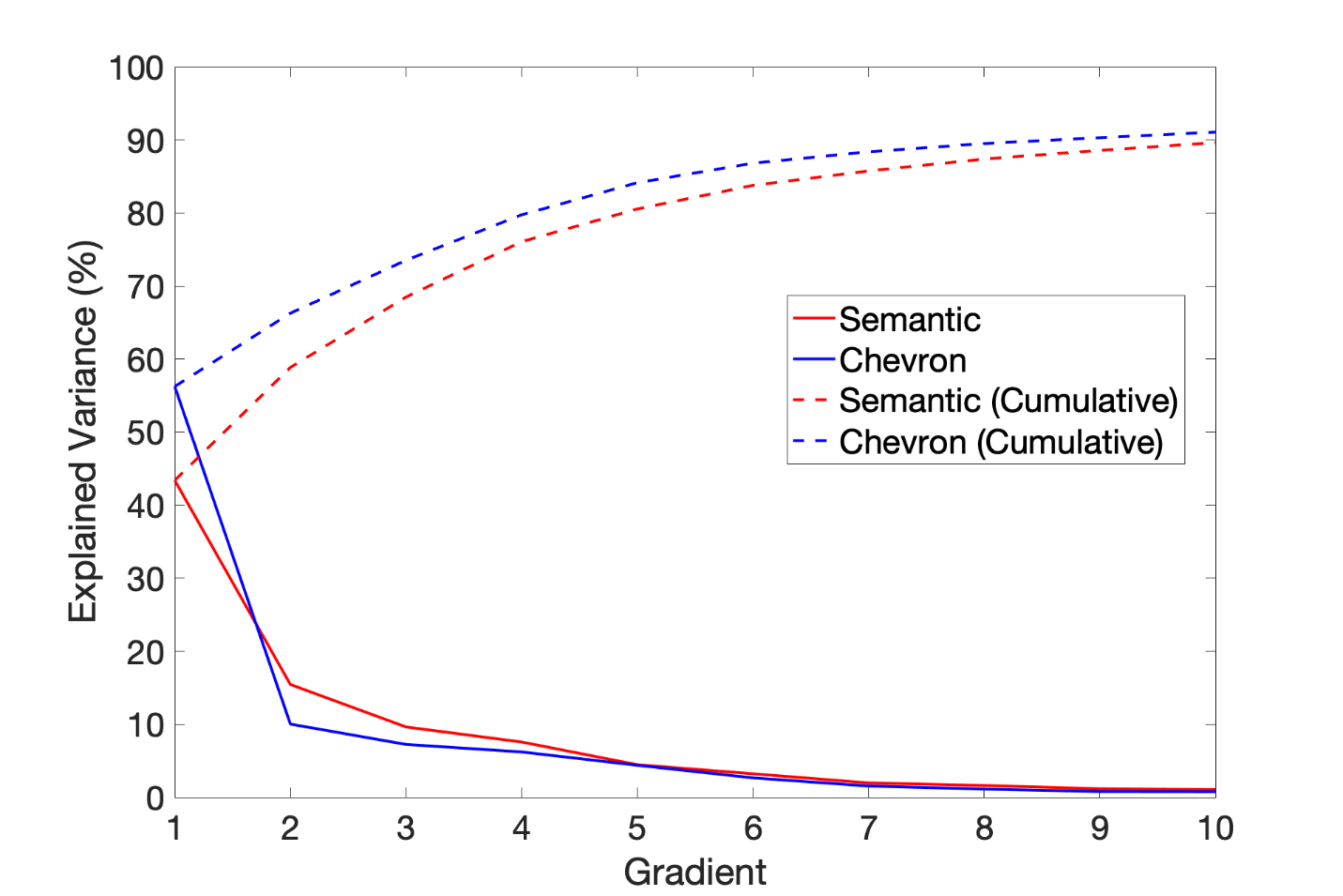

Figure S1. The principal gradient identified by informational connectivity using semantic neural responses and chevron neural responses (upper panel). Variance explained by top ten gradients in each task: dashed line refers to cumulative explained variance; solid line refers to variance explained by each gradient.

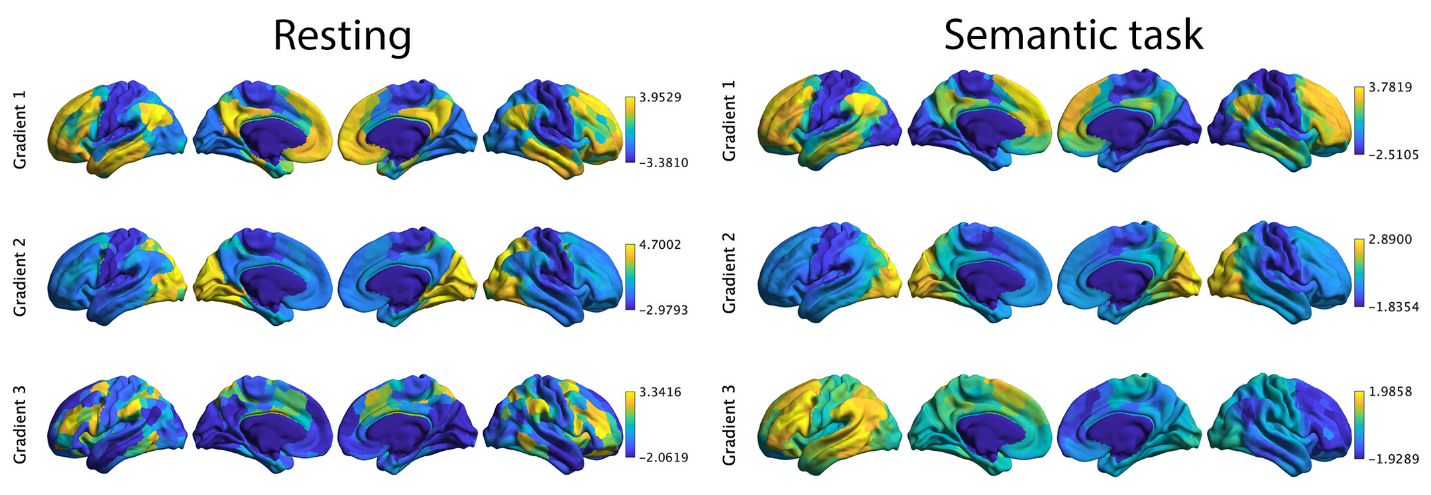

Figure S2. Top three gradients which explained most variance derived from resting connectivity data reported by Margulies et al. (2016) (left panel) and from informational connectivity data from a semantic task (right panel). There were significant correlations for corresponding gradients across these methods. Gradients 1 and 2 showed strong positive correlations, suggesting these dimensions of connectivity are present both during rest and semantic judgements: gradient 1: r = 0.871, Pspin < 0.001; gradient 2: r = 0.425, Pspin < 0.001. In contrast, gradient 3 showed a negative correlation: r = -0.05, Pspin < 0.001.

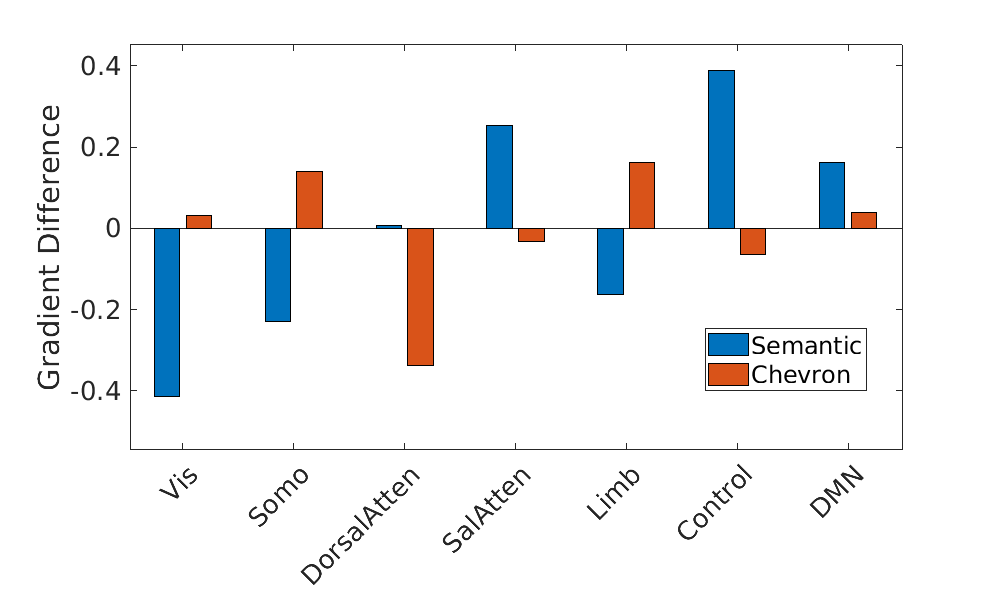

Figure S3. Gradient 1 differences between strong and weak associations in Yeo et al. (2011) 7 networks for semantic and chevron task separately. Vis: visual; Somo: sensorimotor; DorsalAtten: dorsal attention; SalAtten: salience/ventral attention; Limb: limbic; control: frontoparietal; DMN: default mode network.

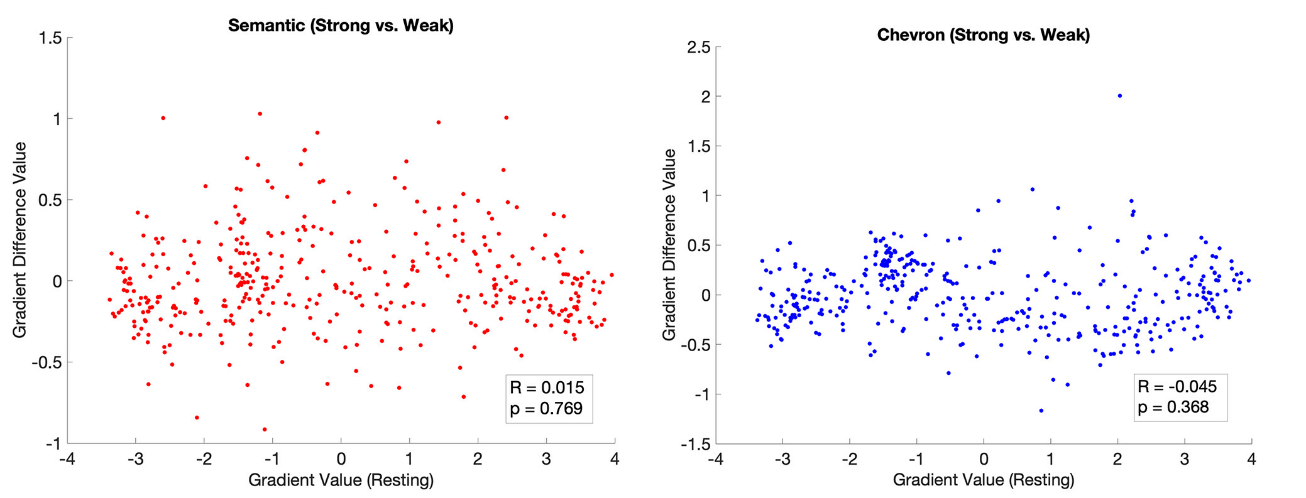

Figure S4. The principal gradient estimated by using traditional functional connectivity (Pearson correlation of averaged time series between any pair of ROIs) was insensitive to the association strength in the semantic task. This principal gradient difference between strong and weak conditions did not show a significant correlation with the principal gradient identified by resting state for both the semantic task (left) and the chevron task (right).

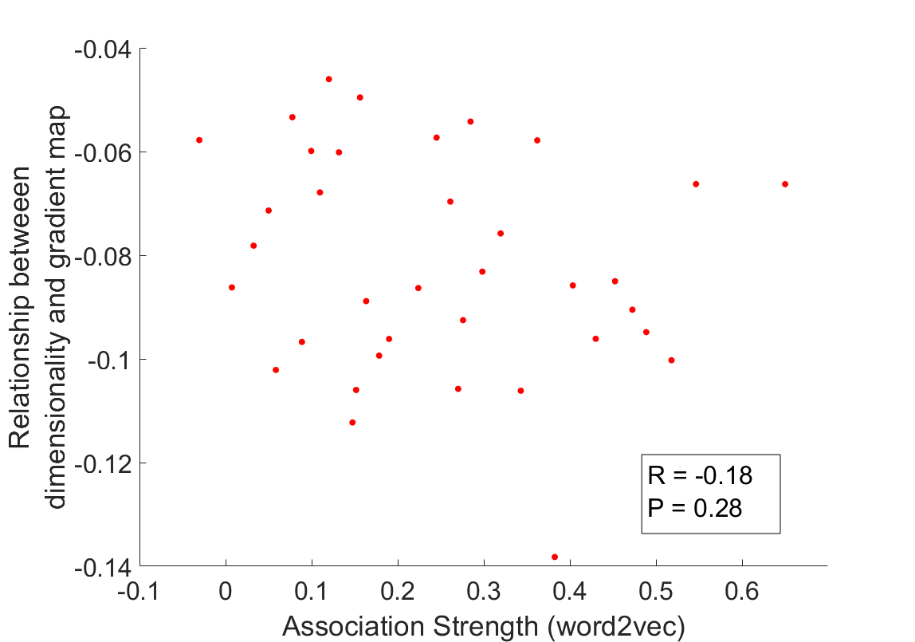

Figure S5. We averaged the dimensionality map in the chevron task for every four trials arranged in order of the word2vec values for the preceding semantic task from weak to strong for this analysis. The relationship between dimensionality and the principal gradient was insensitive to the associative strength of the preceding semantic trial (p = 0.28).

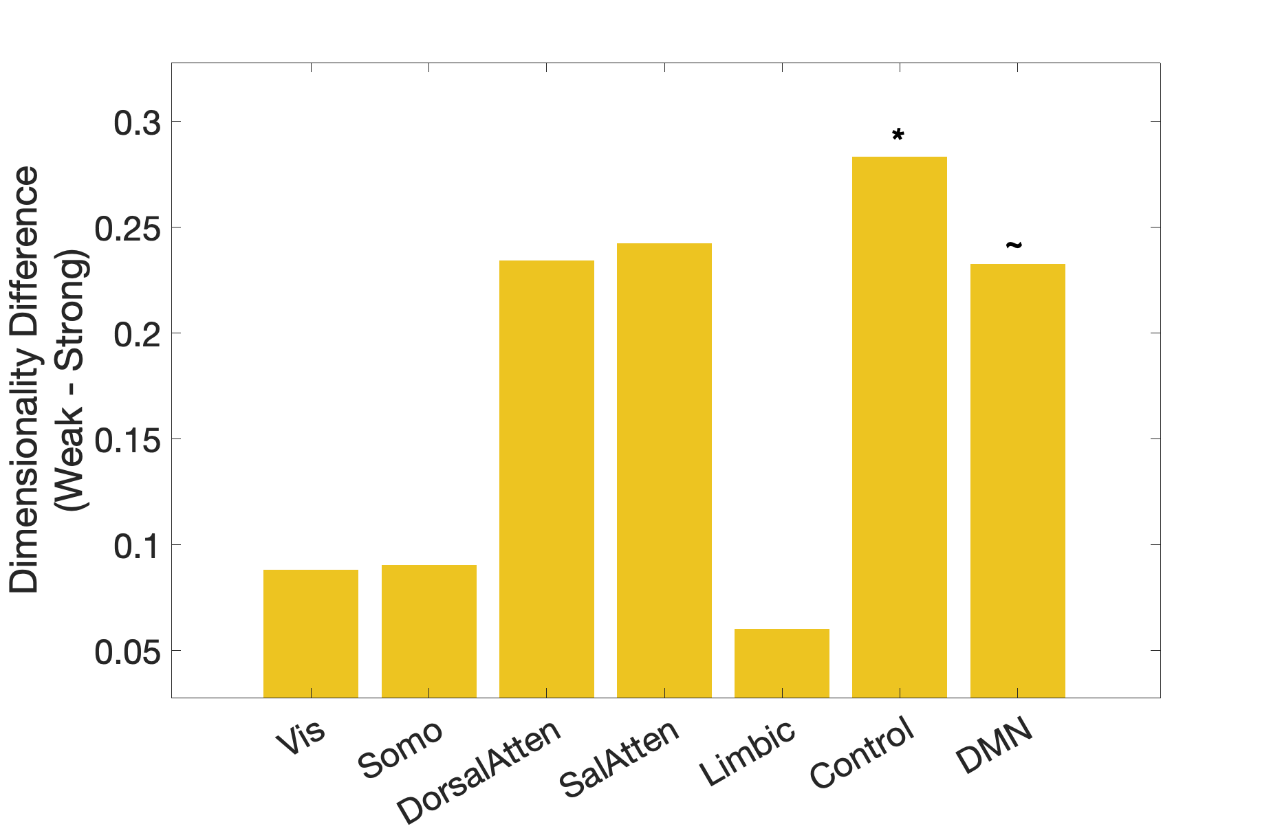

Figure S6. Dimensionality differences between weak and strong association trials in classical resting networks (right) (Yeo et al., 2011).

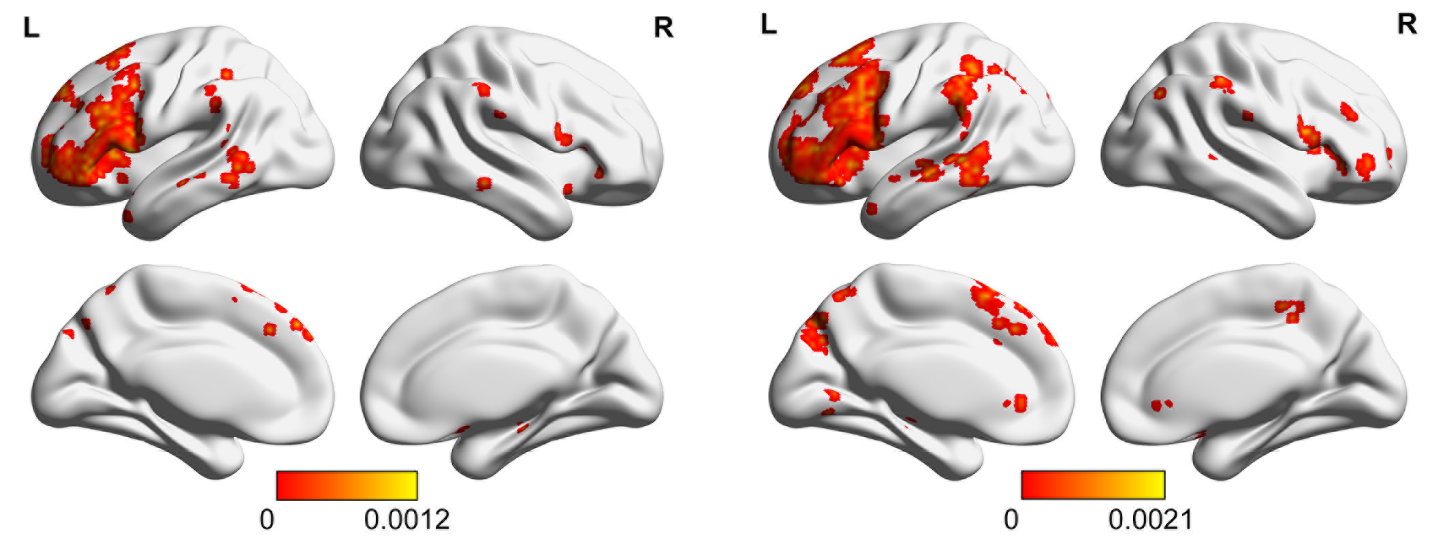

Figure S7. The significance p map of dimensionality differences between weak and strong associations in which the dimensions of the activation pattern were determined using different criteria (explaining 60% (left panel) or 75% (right panel) of the variance), with FDR correction applied, q = 0.05.

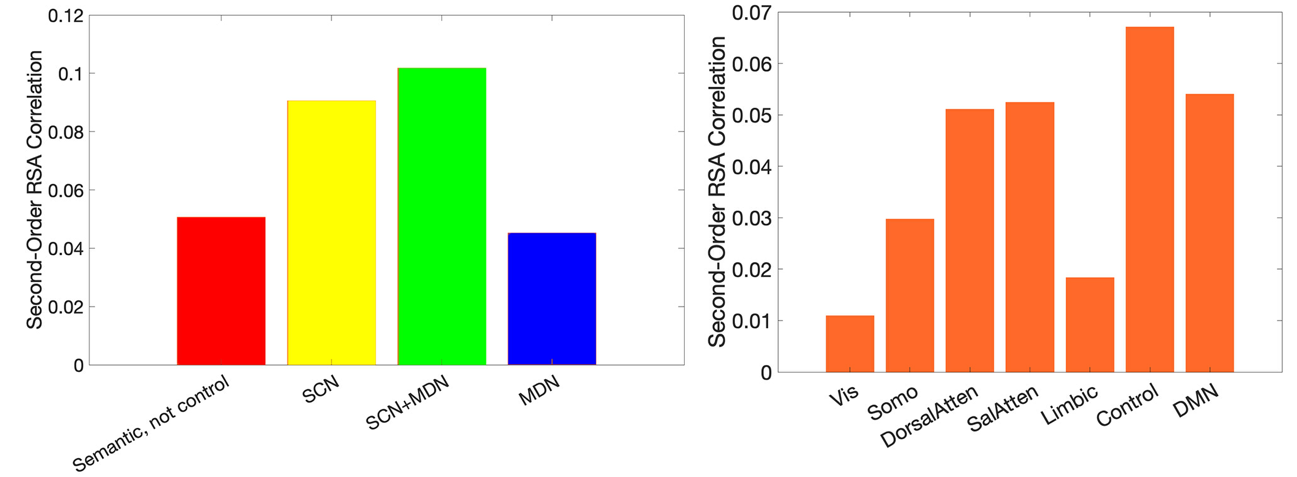

Figure S8. The association between neural and semantic similarity (from ratings of associative strength) estimated using second-order RSA in functional networks implicated in semantic representation and control demands (left) or in resting networks (right) (Yeo et al., 2011). The strongest neural-semantic association effects were found in control-relevant areas in both analyses.

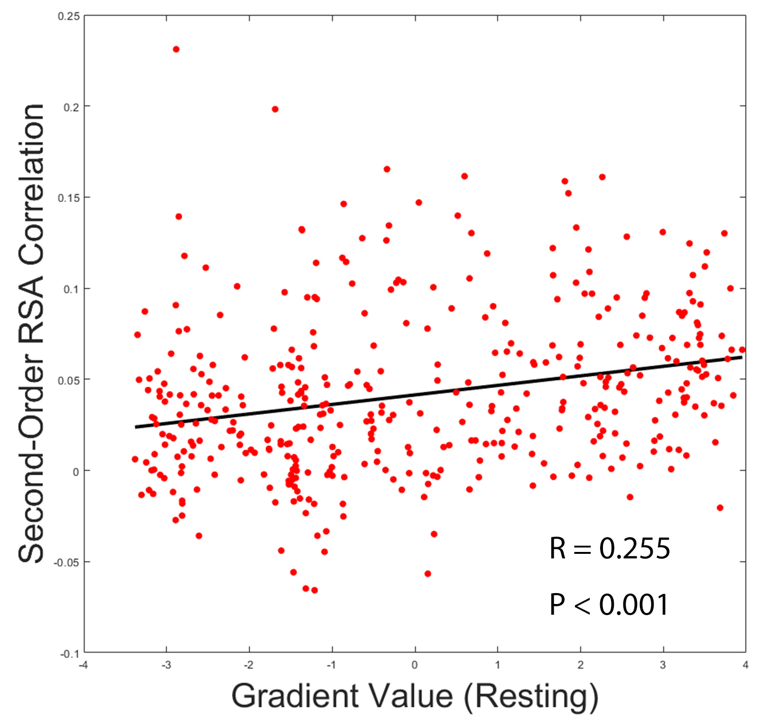

Figure S9. The principal gradient derived from resting data (1) was positively associated with semantic-brain alignment (r = 0.255, p < 0.001).

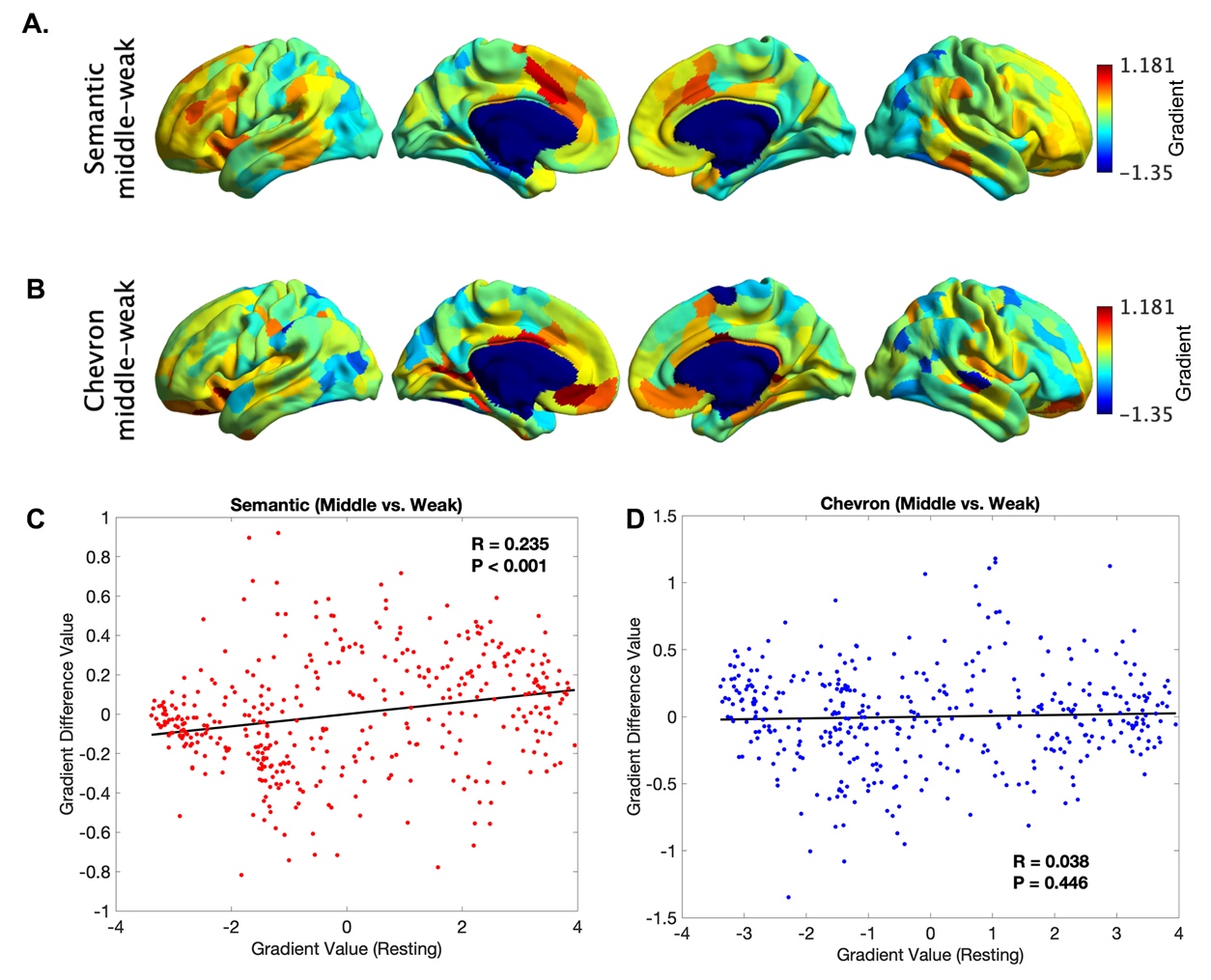

Figure S10. The principal gradient difference between middle 1/3 and bottom 1/3 trials for the semantic neural responses (A) and for chevron task signals (B). The principle gradient difference derived from (A) was positively associated with principle gradient value derived from resting state data (C, r = 0.235, p < 0.001), but no such relationship was found for chevron task (D).

Table S1.

Spin correlation between gradients estimated by traditional connectivity using resting state data and informational connectivity using semantic and chevron neural responses in each hemisphere.

|  |  | Semantic Task | | | | | |
| --- | --- | --- | --- | --- | --- | --- | --- |
|  |  | Left Hemisphere | | | Right Hemisphere | | |
|  |  | Gradient 1 | Gradient 2 | Gradient 3 | Gradient 1 | Gradient 2 | Gradient 3 |
| Resting | Gradient 1 | 0.867*** |  |  | 0.874*** |  |  |
|  | Gradient 2 |  | 0.421*** |  |  | 0.439*** |  |
|  | Gradient 3 |  |  | 0.419*** |  |  | -0.708*** |
|  |  | Chevron Task | | | | | |
|  |  | Left Hemisphere | | | Right Hemisphere | | |
|  |  | Gradient 1 | Gradient 2 | Gradient 3 | Gradient 1 | Gradient 2 | Gradient 3 |
| Resting | Gradient 1 | 0.803*** |  |  | 0.806*** |  |  |
|  | Gradient 2 |  | 0.261*** |  |  | 0.246*** |  |
|  | Gradient 3 |  |  | -0.060*** |  |  | -0.833*** |

Gradients 1 and 2 showed similar spatial patterns when derived from informational connectivity metrics in the semantic and chevron tasks and from resting state data. However, gradient 3 showed a task-specific state which captured the separation of left hemisphere from right hemisphere for the semantic task.

MARGULIES, D. S., GHOSH, S. S., GOULAS, A., FALKIEWICZ, M., HUNTENBURG, J. M., LANGS, G., BEZGIN, G., EICKHOFF, S. B., CASTELLANOS, F. X., PETRIDES, M., JEFFERIES, E. & SMALLWOOD, J. 2016. Situating the default-mode network along a principal gradient of macroscale cortical organization. *Proceedings of the National Academy of Sciences,* 113**,** 12574-12579.

YEO, B. T. T., KRIENEN, F. M., SEPULCRE, J., SABUNCU, M. R., LASHKARI, D., HOLLINSHEAD, M., ROFFMAN, J. L., SMOLLER, J. W., ZÖLLEI, L., POLIMENI, J. R., FISCHL, B., LIU, H. & BUCKNER, R. L. 2011. The organization of the human cerebral cortex estimated by intrinsic functional connectivity. *Journal of Neurophysiology,* 106**,** 1125-1165.
